# Impact of hydrostatic pressure on organic carbon cycling of the deep-sea microbiome

**DOI:** 10.1101/2022.03.31.486587

**Authors:** Chie Amano, Zihao Zhao, Eva Sintes, Thomas Reinthaler, Julia Stefanschitz, Murat Kisadur, Motoo Utsumi, Gerhard J. Herndl

**Affiliations:** Department of Functional and Evolutionary Ecology, University of Vienna, Djerassiplatz 1, 1030 Vienna, Austria; Instituto Español de Oceanografía-CSIC, Centro Oceanográfico de Baleares, Muelle de Poniente s/n, Apdo 291, 07015 Palma de Mallorca, Spain; Faculty of Life and Environmental Sciences, University of Tsukuba, Tennodai 1-1-1, Tsukuba, Ibaraki 305-8572, Japan; NIOZ, Department of Marine Microbiology and Biogeochemistry, Royal Netherlands Institute for Sea Research, Utrecht University, 1790 AB Den Burg, Texel, The Netherlands; Vienna Metabolomics Center, University of Vienna, Djerassiplatz 1, A-1030 Vienna, Austria

## Abstract

Deep-sea microbial communities are exposed to high hydrostatic pressure. While some of these deep-sea prokaryotes are adapted to high-pressure conditions, the contribution of piezophilic (i.e., pressure-loving) and piezotolerant prokaryotes to the total deep-sea prokaryotic community remains unknown. Here we show that the metabolic activity of prokaryotic communities is increasingly inhibited with increasing hydrostatic pressure. At 4,000 m depth, the bulk heterotrophic prokaryotic activity under *in sit*u hydrostatic pressure was only about one-third of that measured on the same community at atmospheric pressure conditions. Only ∼5% of the bathypelagic prokaryotic community are piezophilic while ∼85% of the deep-sea prokaryotes are piezotolerant. A small fraction (∼10%) of the deep-sea prokaryotes is piezosensitive (mainly members of Bacteroidetes, Alteromonas) exhibiting specific survival strategies at meso- and bathypelagic depths. These piezosensitive bacteria elevated their activity by more than 100-fold upon depressurization. Hence, the consistently higher bulk metabolic activity of the deep-sea prokaryotic community measured upon depressurization is due to a rather small fraction of the prokaryotic community. Overall, the heterotrophic prokaryotic activity in the deep-sea is substantially lower than hitherto assumed with major impacts on the oceanic carbon cycling.

---

The water column of the deep-sea is a dark and typically cold realm (0–4°C) with hydrostatic pressure increasing with depth. Prokaryotic abundance and activity are also decreasing with depth, generally interpreted as a reflection of decreasing energy supply rates with depth^1^. After the submersible *Alvin* accidently sank almost 50 years ago, Jannasch *et al*. found that food left in *Alvin* at 1,540 m depth for more than ten months was remarkably well-preserved^2^. They concluded that the high hydrostatic pressure prevented deep-sea microbes from utilizing this food source. Subsequently, studies on the effect of hydrostatic pressure on deep-sea prokaryotes were performed^3^, however, revealed inconclusive results. Most of these studies applied relatively high substrate concentrations to determine metabolic rates^2^ or were performed in the Mediterranean Sea with its exceptionally high temperature of the bathypelagic waters (∼13°C), which can influence the metabolism and physiology of deep-sea microbes^4-6^. Due to the methodological difficulties to measure prokaryotic activity under *in situ* pressure conditions, only a few comparative measurements of prokaryotic activity under *in situ* pressure and depressurized conditions are available from the meso- and bathypelagic global ocean despite the potential impact hydrostatic pressure might have on deep-sea microbial activity and on understanding the ocean biogeochemical cycle^3,7-9^.

While the activity of sea-surface microbial communities is reduced or inhibited by hydrostatic pressure at about 10 MPa (corresponding to a depth of 1,000 m)^10^, some deep-sea microbes exhibit a piezophilic (i.e., optimal growth at pressures >0.1 MPa) and piezotolerant lifestyle with specific adaptions to high hydrostatic pressure, low temperature and low nutrient conditions^11^. Comparing genomes from obligate piezophilic and piezosensitive microbes grown under low temperature (optimal growth of the piezophiles at 6–10°C) indicated an adaptation to high hydrostatic pressure in piezophiles in membrane fluidity, stress response and cell motility^12^, consistent with previous culture based studies^11^.

Commonly, the prokaryotic heterotrophic carbon demand (PCD) of deep-sea microbes is calculated from heterotrophic biomass production and respiration measurements based on ship-board incubations under atmospheric pressure conditions, assuming that pressure changes do not affect metabolic rates. Estimates of the PCD in the meso- and bathypelagic layers of the Atlantic revealed that the PCD is about one order of magnitude higher than the supply of particulate organic carbon via sinking particles^13^. A similar conclusion was reached for the Pacific albeit using a different approach^14^. This mismatch between the PCD and particulate organic carbon supply via sinking particles indicates some fundamental errors in our estimates on deep-sea prokaryotic activity and/or on the magnitude of sinking organic matter flux^1,9,15,16^.

## Heterotrophic microbial activity at *in situ* pressure

The heterotrophic prokaryotic activity was determined under *in situ* pressure conditions throughout the water column down to bathypelagic layers in the Atlantic, Pacific and Indian sector of the Southern Ocean (Extended Data Fig. 1 and Extended Data Table 1). Heterotrophic prokaryotic activity was assessed via the incorporation of radiolabeled leucine into proteins^17^ using the *In Situ* Microbial Incubator (ISMI; Extended Data Fig. 2, manufactured by NiGK Corporation, Japan). The ISMI collects and incubates water at depths down to 4,000 m with substrate added such as ^3^H-leucine at the depth of sampling. Thus, the ISMI allows determining prokaryotic activity without changes of the hydrostatic pressure and temperature, hence under *in situ* conditions (see Methods). The results obtained from these *in situ* incubations using the ISMI were compared to measurements on samples collected at the same site and depth as those of ISMI but under atmospheric pressure on board of the respective research vessel. Care was taken to prevent any contamination with organic and inorganic matter in all incubation bottles and the incubation temperature was the same as the temperature in the *in situ* incubations (Methods and Extended Data Table 2).

Generally, heterotrophic prokaryotic activity decreased with depth, however, under *in situ* pressure more than under atmospheric pressure conditions (*P* = 0.048 for the slopes of log–log fits assuming power law function; Extended Data Fig. 3). For samples collected at 500 m depth, the impact of hydrostatic pressure was small reaching about 75 ± 10% (mean ± s.d., *n* = 4) of the activity measured at atmospheric pressure (Fig. 1 and Extended Data Table 2). The difference in prokaryotic activity between *in situ* and atmospheric pressure conditions was most pronounced in the bathypelagic waters. *In situ* prokaryotic activity at ∼1,000 m depth was ∼60 ± 10% (mean ± s.d., *n* = 3) of that under atmospheric pressure. At the base of the bathypelagic waters (∼4,000 m depth) *in situ* prokaryotic activity was only ∼30 ± 15% (mean ± s.d., *n* = 4) of that measured under atmospheric pressure (Fig. 1 and Extended Data Table 2). Thus, bulk heterotrophic prokaryotic activity is greatly reduced in the bathypelagic realm under *in situ* pressure conditions. The question whether most of the members of the microbial community are suppressed in their metabolic activity or only a small fraction responding to depressurization with elevated activity was addressed using single-cell activity measurements.

**Figure 1.**
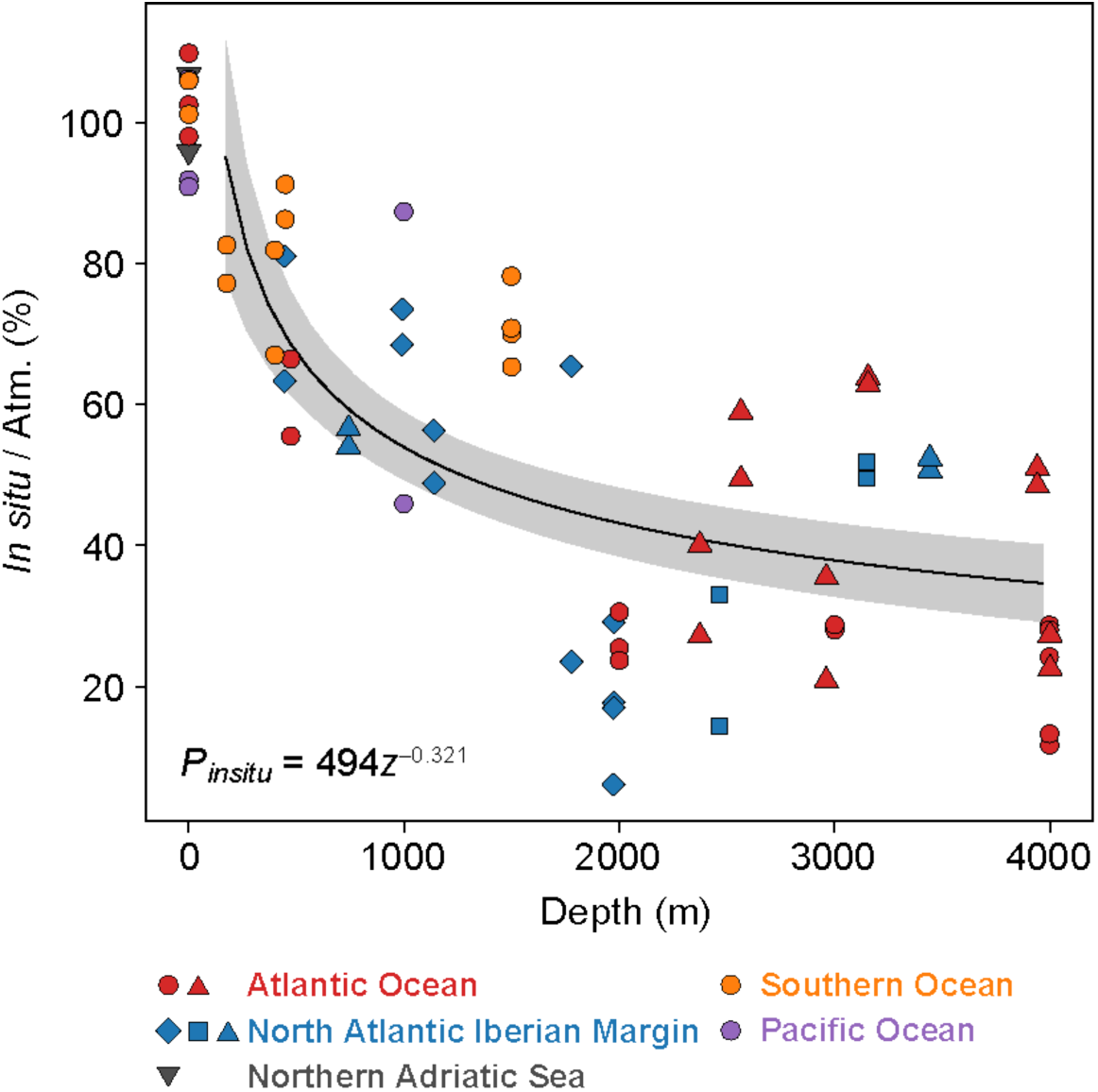
Percentage of bulk leucine incorporation rates obtained *in situ* normalized to those at atmospheric pressure conditions. Symbols correspond to the different research expeditions (Extended Data Fig. 1). Regression equation is a power law function: *P*_*insitu*_ = 494*z*^−0.321^ (*n* = 56) where *P*_*insitu*_ is the percentage of *in situ* leucine incorporation rate normalized to leucine incorporation rate under atmospheric pressure (Atm.) and *z* is depth (in m). Shaded area indicates 95% confidence interval. Note that the data points at 0 m (*n* = 4) correspond to instrumental tests in which epi- to bathypelagic waters were incubated with the ISMI under atmospheric pressure conditions and compared to bottle incubations used for atmospheric pressure incubations to assess the potential bias associated to the instrument. These points are excluded from calculating the regression line.

## Leucine incorporation rates at a single-cell level

Single-cell prokaryotic activity under *in situ* and atmospheric pressure conditions was determined on three mesopelagic (∼400–750 m depth) and six bathypelagic samples (∼1,500– 4,000 m depth) collected in the Atlantic and Southern Ocean using microautoradiography with ^3^H labeled leucine combined with catalyzed reporter deposition fluorescence *in situ* hybridization (Methods). Using microautoradiography, the silver grain halo around single cells indicating uptake of radiolabeled leucine serves as a proxy for single-cell prokaryotic activity^18,19^ (see Methods). There was no detectable difference between *in situ* and on-board incubations at atmospheric pressure conditions in prokaryotic abundance (paired *t*-test, *P* = 0.724; Extended Data Table 3) and in the abundance of cells taking up leucine (paired *t*-test, *P* = 0.905). However, the total size of the silver grain halo around the cells taking up leucine, i.e., cell-specfic leucine uptake was higher under atmospheric pressure than under *in situ* hydrostatic pressure conditions (Extended Data Table 3). This is in agreement with the higher bulk leucine incorporation rates obtained under atmospheric than under *in situ* pressure conditions (Extended Data Table 2). Highly active cells (>0.5 amol leucine uptake cell^−1^ day^− 1^) were found in the samples incubated under atmospheric pressure, hence depressurized conditions (Fig. 2a, b). These cells were generally low in abundance (1–5% of total cells taking up leucine) and were essentially absent in the samples where *in situ* pressure was maintained (Extended Data Table 3). This highly active fraction detected under depressurized conditions can be considered as the piezosensitive fraction of the prokaryotic community. Apparently, relieving these piezosensitive prokaryotes from hydrostatic pressure provoked the increase of bulk leucine incorporation rates. Analysing the changes of cell-specific uptake rates from *in situ* to atmospheric pressure conditions over the whole activity range allows estimating the abundance of piezosensitive, piezotolerant and piezophilic prokaryotes. In the bathypelagic waters, 1–30% of cells taking up leucine were classified as piezosensitives (Extended Data Table 3). The majority (≥80%) of the deep-sea prokaryotes, however, was piezotolerant (Extended Data Table 3, except one sample). Only a small fraction (∼5%) appeared to be piezophilic exhibiting higher cell-specific activity under *in situ* pressure than under depressurized conditions (Extended Data Table 3). Only in one sample from 4,000 m depth, ∼20% of the cells were considered piezophilic (Extended Data Table 3). Leucine uptake rates of the piezophiles were generally low and never exceeded that of the piezosensitive fraction.

**Figure 2.**
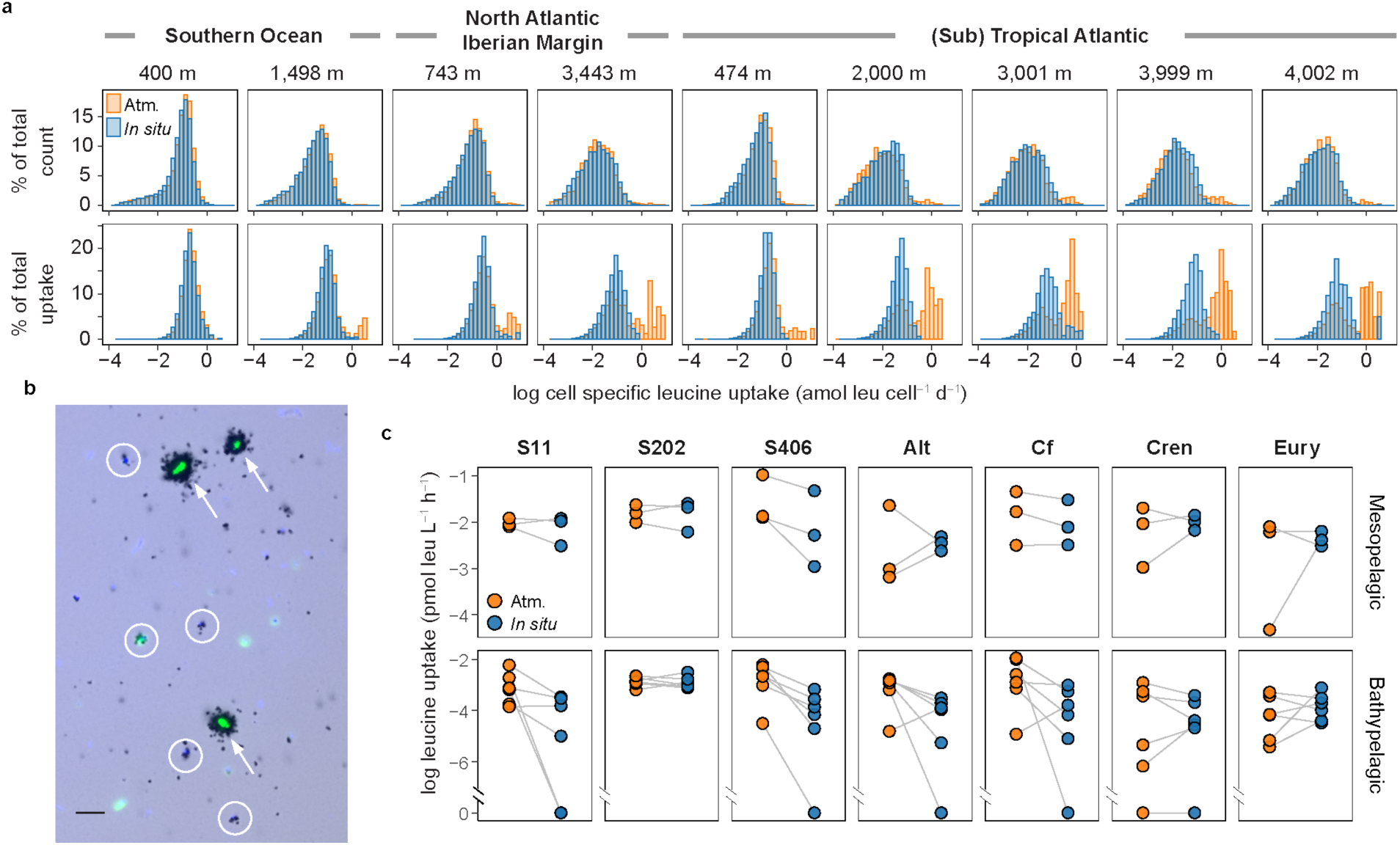
Cell-specific leucine uptake by prokaryotes. **a**, Distribution of cell-specific leucine uptake expressed as the percentage of total active cell-counts (upper panels) and the percentage of total uptake (lower panels). Water was collected at meso- and bathypelagic depths and incubated under *in situ* and atmospheric pressure (Atm.) conditions (Extended Data Tables 1, 2). **b**, A microscopic view of a bathypelagic sample (2,000 m) collected in the Atlantic and incubated under atmospheric pressure condition. Black halos around the cells are silver grains corresponding to their activities. The highly active cells (>0.5 amol leu cell^−1^ d^−1^, indicated by arrows) were barely found in *in situ* pressure incubations. Typical low-activity cells in the bathypelagic depths are indicated by circles. Green fluorescence, EUB338 probe mix; light blue, DAPI-stained cells. Scale bar; 5 µm. **c**, Leucine uptake by taxonomical groups: S11, SAR11 clade; S202, SAR202 clade; S406, SAR406 clade; Alt, Alteromonas; Cf, Bacteroidetes; Cren, Thaumarchaeota; Eury, Euryarchaeota. The gray line connects the same location and depth between *in situ* and Atm. samples representing the change in leucine uptake beween the two incubation conditions.

Significantly higher heterotrophic activity upon depressurization, hence a piezosensitive response, was observed for several members of the bacterial community, particularly in Bacteroidetes (paired *t*-test, *n* = 18, *P* = 0.025) and SAR406 (Marinomicrobia; Wilcoxon signed-rank test, *n* = 18, *P* = 0.004), and Alteromonas especially from bathypelagic waters (paired *t*-test, *n* = 12, *P* = 0.012) (Fig. 2c and Extended Data Fig. 4a). In contrast to these piezosensitive prokaryotes, SAR202 (Chloroflexi) showed no significant differences in leucine uptake between *in situ* and atmospheric pressure conditions (Wilcoxon signed-rank test, *n* = 18, *P* = 0.734 and *P* = 0.496 for SAR202 leucine uptake rates and relative abundance of SAR202 taking up leucine, respectively; Fig. 2c and Extended Data Fig. 4b), indicative of a piezotolerant lifestyle. Thaumarchaeota contributed ∼10% to the total prokaryotic abundance (Extended Data Fig. 5). However, only small fraction of Thaumarchaeota in the bathypelagic waters (∼10% of thaumarchaeal cells) took up leucine under both *in situ* and atmospheric pressure conditions (Extended Data Fig. 4b). No difference in leucine uptake rates in Thaumarchaeota under *in situ* and atmospheric pressure was observed (Wilcoxon signed-rank test for leucine uptake, *n* = 18, *P* = 0.834; paired *t*-test for relative abundance of cells taking up leucine, *n* = 18, *P* = 0.148; Fig. 2c and Extended Data Fig. 4b), indicating that bathypelagic Thaumarchaeota are probably piezotolerant.

Taken together, we conclude that the vast majority of the deep-sea prokaryotic community is piezotolerant and only minor fractions are piezosensitive and piezophilic. While members of the Bacteroidetes and Alteromonas such as the genus Colwellia have been shown to be piezophilic^20,21^, we consistently found that both Bacteroidetes and Alteromonas are piezosensitive. This might indicate that most members of both taxa originate from surface waters and are associated with particles sinking out of the surface layers into the ocean’s interior. To obtain a better insight into the metabolic response of deep-sea prokaryotes upon depressurization the metaproteome of abundant piezosensitive and piezotolerant bacterial taxa was analyzed.

## Depth related changes in the metaproteome

While protein expression and function are influenced by hydrostatic pressure, monomeric proteins are rather stable compounds under a moderate pressure range (<400 MPa)^22^. Protein synthesis requires a certain period of time via transcription and translation and is related to the growth rate of heterotrophic prokaryotes in the ocean^23^. Due to the generally low growth rates of the heterotrophic prokaryotic community in the deep-sea^13^, proteins extracted from deep-sea prokaryotes were likely expressed under *in situ* conditions. We performed metaproteomic analyses with a focus on Alteromonas, Bacteroidetes and SAR202 since single-cell activity measurements indicated that the former two bacterial taxa are piezosensitive while SAR202 is piezotolerant (Fig. 2c and Extended Data Fig. 4). We aimed at deciphering strategies of these different taxa to adapt to hydrostatic pressure. Based on gene ontology (GO)^24^, there is apparently no universal adaptation among these three bacterial taxa related to changes in hydrostatic pressure (Fig. 3a). However, taxa-specific differences were detectable related to the sampling depth and hence, hydrostatic pressure (Fig. 3b and Supplementary Tables 1 and 2).

**Figure 3.**
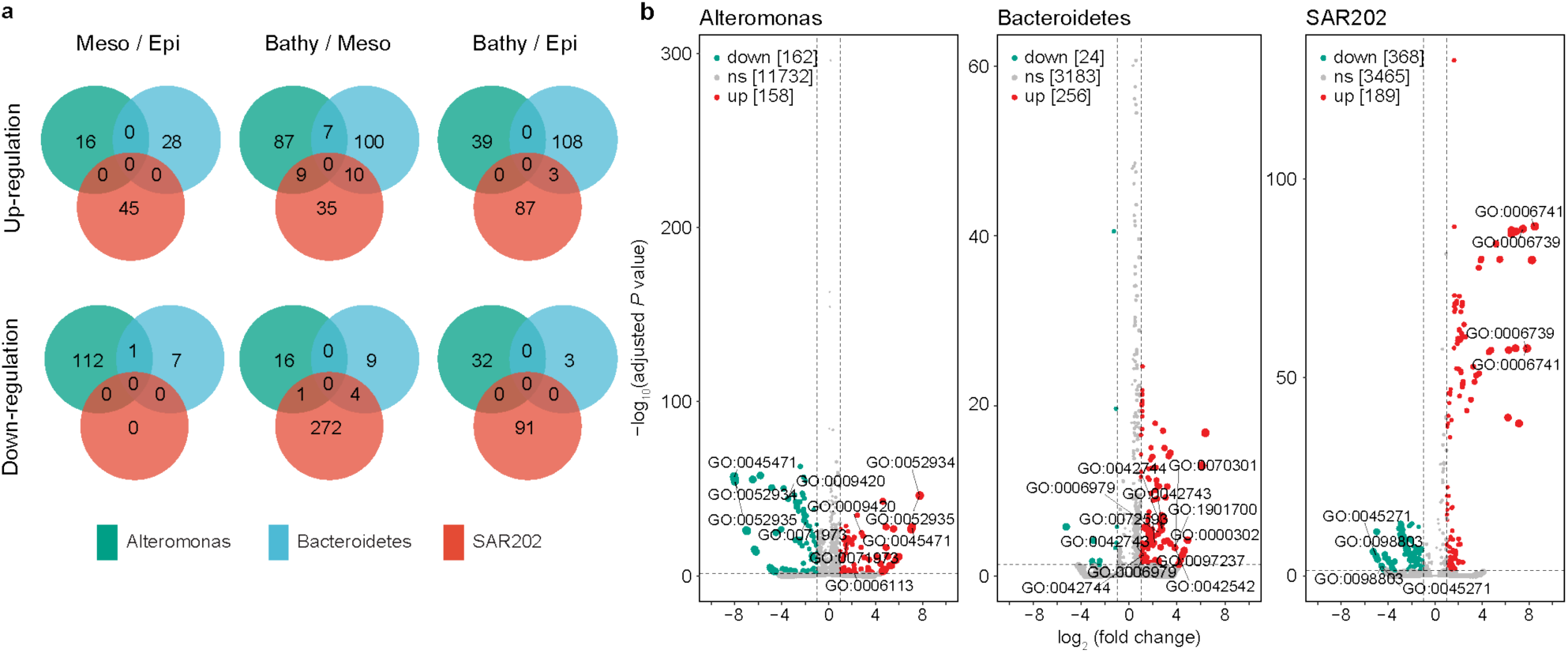
Depth related changes in the metaproteome of three abundant deep-sea bacterial taxa. **a**, Venn diagrams indication the number of shared and unique up- and down-regulated proteins among Alteromonas, Bacteroidetes and SAR202 of meso-*versus* epipelagic layers, bathy-*versus* mesopelagic layers and bathy-*versus* epipelagic layers. Numbers indicate the protein abundance. Epi: epipelagic, Meso: mesopelagic, Bathy: bathypelagic waters. **b**, Comparison of expressed proteins produced by Alteromonas, Bacteroidetes and SAR202. Significance of the change between depth layers is indicated by different colours: not significant (ns), *P* ≥ 0.05; up regulated proteins (up), *P* < 0.05 and log_2_ fold change ≥1; down regulated proteins (down), *P* < 0.05 and log_2_ fold change ≤ –1.

Bacteroidetes up-regulated the response to oxidative stress (i.e., response to hydrogen peroxide, reactive oxygen species and oxygen-containing compounds; Supplementary Table 1) in the bathypelagic as compared to the epipelagic layer. Culture based analyses revealed that resistance against oxidative stress in deep-sea prokaryotes is an adaptation to high hydrostatic pressure and low temperature^25^. Particle-associated Bacteroidetes were suggested to exhibit a piezosensitive lifestyle^26^. Also Alteromonas living in the bathypelagic realm exhibit a pronounced tendency towards a particle-associated lifestyle^27^. This particle-associated lifestyle of Altermonas in the bathypelagic layers allows them to access organic matter at higher concentrations on particles than in the ambient water^28^. Moreover, particles, such as deep-sea marine snow^29^ provide a micro-environment potentially favouring fermentation (i.e., anaerobic respiration). Alteromonas showed flexibility to the change of hydrostatic pressure by down- and up-regulating the same genes depending on the depth layers (e.g., GO:0045471, GO:0052934, GO:0052935, GO:0009420, GO:0071973; Fig. 3b, Supplementary Tables 1, 2). In the bathypelagic compared to the mesopelagic layers, the flagellum synthesis (GO:0009420, GO:0071973; Supplementary Table 1) and the fermentation pathway were up-regulated in Alteromonas. The response to ethanol (GO:0045471; Supplementary Table 1) was also up-regulated in the bathypelagic realm probably related to the fermentation of algal derived polysaccharides. In addition, the up-regulation of alcohol dehydrogenase activity (GO:0052934, GO:0052935) suggests counteracting oxidative damage due to high pressure and low temperature in bathypelagic Alteromonas.

In SAR202, NADP biosynthesis and metabolism (GO:0006741, GO:0006739) were up-regulated in the bathypelagic realm while the respiratory chain complex 1 (GO:0045271, GO:0098803) was five-fold down-regulated compared to the mesopelagic realm (Supplementary Tables 1 and 2). Respiratory chains are known to be affected by hydrostatic pressure, including in piezophilic bacteria^30^. This might be interpreted as an adaptation to the limited substrate availability in bathypelagic compared to the mesopelagic waters. It might be a strategy allowing for a similar activity under *in situ* and depressurized conditions as revealed by single-cell analysis (Extended Data Fig. 4a).

Taken together, there are taxa-specific modifications in Alteromonas and Bacteroidetes at the proteome level in bathypelagic cells resulting probably in a higher energy expenditure to maintain a certain level of metabolism under high hydrostatic pressure as described in a previous study^31^. Upon depressurization, these specific adaptations are not required leading overall, to a higher metabolic activity of Alteromonas and Bacteroidetes under atmospheric pressure than at deep-sea pressure conditions. In contrast, SAR202 as a representative of the vast majority of piezotolerant prokaryotes down-regulate the respiratory complex 1 and up-regulate NADP biosynthesis in the bathypelagic realm to maintain the metabolic activity level under contrasting hydrostatic pressure condition.

## Vertical transport of prokaryotes through the water column

Our results from single-cell analyses and metaproteomics support the conclusion that the piezosensitive microbes (mainly Alteromonas and Bacteroidetes) most likely originated from the upper water column. These piezosensitive Bacteria instantly responded to the depressurization within the relatively short incubation period required to measure heterotrophic activity (3–12 h). The abundance of piezosensitive prokaryotes decreased exponentially with depth (Extended Data Fig. 6). Occasionally, the fraction of piezosensitive cells of the total active community was high (20–30%; Extended Data Table 3), tentatively indicating episodically rapid transport of these cells on sinking particles such as marine snow. Alteromonas and Bacteroidetes are known to be ubiquitous, generalistic/opportunistic bacterial taxa; the former are capable of rapidly exploiting available substrate^28^ and are abundant in marine snow from euphotic to bathypelagic waters^32^. Bacteroidetes are abundant in particle-rich epipelagic waters utilizing preferentially high molecular weight organic matter associated to phytoplankton blooms^28,33-35^ and are found on sinking particles at bathypelagic depth during elevated particle export events^36^. Hence, these bacterial taxa are likely transported from the surface to the deep waters via sedimenting particles.

The stimulation of heterotrophic prokaryotic activity under atmospheric pressure conditions could be caused by the release of intracellular organic matter from organisms upon depressurization. If we assume a prokaryotic carbon content^37,38^ of 10 fg C cell^−1^, the mean prokaryotic cell abundance in the bathypelagic waters (2.9 ± 1.4 × 10^4^ cells mL^−1^, *n* = 4; Extended Data Table 3) would result in 0.29 ± 0.14 μg C biomass L^−1^. The difference of the bulk heterotrophic bacterial biomass production between *in situ* incubations and under atmospheric pressure conditions was 0.003–0.029 pmol Leu L^−1^ h^−1^. Using a conversion factor of 1.55 kg C biomass mol^−1^ leucine incorporated which is at the high end of conversion factors for deep-sea heterotrophic prokaryotes^39^ and a growth yield of 50%, this translates into an additional organic carbon demand of 0.43 ± 0.40 ng C L^-1^ (mean ± s.d., *n* = 4) under atmospheric as compared to *in situ* conditions. This is equivalent to 0.1–0.4% of the bathypelagic prokaryotic biomass. Thus, if only a few prokaryotic cells would burst during the depressurization, it would be sufficient to stimulate heterotrophic prokaryotic activity under atmospheric pressure. No signs of cell debris, however, were noticed in microscopic examinations.

Other parameters potentially being altered upon depressurization are oxygen and carbon dioxide concentrations. Oxygen availability at all our study sites was not a growth limiting factor for aerobic prokaryotes (Extended Data Table 1) nor the changes in CO_2_ concentrations and the associated small pH changes upon depressurization.

Regardless of whether or not some deep-sea prokaryotes released organic matter into the water or some physico-chemical parameters changed upon depressurization and thus provoked the higher metabolic activity of the bulk deep-sea heterotrophic prokaryotic community under atmospheric pressure, the major conclusion of our study remains: measuring deep-sea prokaryotic activity under atmospheric pressure conditions leads to a substantial overestimation of the actual *in situ* bulk prokaryotic activity. Consequently, deep-sea prokaryotic activity should be determined by maintaining the *in situ* hydrostatic pressure conditions to allow better constraining the deep-sea carbon flux ^20,40,41^ as heterotrophic prokaryotes are by far the most important remineralizers of organic carbon in the ocean.

## Implication on the deep-sea carbon budget

Apparently, heterotrophic biomass production of deep-sea prokaryotes has been overestimated in the past, since almost all the estimates have been based on measurements performed under atmospheric pressure conditions^1,9^. It is likely that the biomass production and respiration of the bulk prokaryotic community are reduced proportionally under *in situ* pressure conditions. Hence, the growth efficiency remains probably unaffected under *in situ* pressure conditions. The heterotrophic prokaryotic carbon demand (PCD, sum of carbon biomass production and respiration) at several depth horizons of the ocean water column can be compared to the estimated particle flux into the ocean’s interior using heterotrophic prokaryotic production (see Methods). Assuming a growth efficiency of 8% and 3% for meso- and bathypelagic layers, respectively^13,42^, and applying the leucine to carbon conversion factors of 1.55 and 0.44 kg C mol^−1^ leucine incorporated^42,43^, the estimated PCD obtained from *in situ* activity measurements and the particulate organic carbon (POC) supply is largely balanced (Extended Data Fig. 7). A conversion factor of 0.44 kg C mol^−1^ leucine incorporated has been reported for heterotrophic mesopelagic prokaryotic communities^39,44^. Hence, it is likely that this conservative conversion factor is reflecting closely the actual PCD in the bathypelagic realm.

Our study shows that the bulk prokaryotic heterotrophic activity in the deep-sea is substantially inhibited by the hydrostatic pressure in the meso- and bathypelagic realm of the ocean. Thus, despite the fact that the prokaryotic community composition is depth stratified^45^, the small fraction (∼10%) of piezosensitive prokaryotes transported into the deep ocean via particle sedimentation can strongly affect bathypelagic heterotrophic prokaryotic activity measurements performed under atmospheric pressure conditions. Also, only a rather minor fraction (about 5%) appears to be piezophilic in the bathypelagic ocean.

Overall, by taking the inhibitory effect of hydrostatic pressure on the metabolism of the bulk deep-sea heterotrophic prokaryotic community into consideration, the heterotrophic prokaryotic carbon demand and particulate organic carbon supply appears to be largely balanced in the global ocean’s interior. Hence the reported mismatch between organic carbon supply and prokaryotic carbon demand in the bathypelagic realm is largely due to an overestimation of the heterotrophic prokaryotic activity when measured under atmospheric pressure conditions.

## Methods

### Collecting and incubating samples at *in situ* hydrostatic pressure and under atmospheric conditions

For measuring heterotrophic biomass production under *in situ* hydrostatic pressure conditions, water samples were collected and incubated with the autonomous *In Situ* Microbial Incubator (ISMI, NiGK corporation, Japan; Extended Data Fig 2). The ISMI is lowered via a winch from the research vessel to the pre-defined depth. The ISMI is a programmable device consisting of 500 mL polycarbonate incubation and fixation bottles and peristaltic pumps (∼150 mL min^−1^). These parts are connected by silicone tubing. All parts in direct contact with the water samples were thoroughly cleaned. After lowering the ISMI to the pre-defined depth, ambient seawater was pumped by the peristaltic pump into duplicate or triplicate incubation bottles to which ^3^H-leucine (5 nM final concentration, 10 nM for epipelagic samples of the Southern Ocean [3, 4, 5-^3^H] L-leucine with a specific activity ranging between 110–120 Ci mmol^−1^, either from BioTrend or PerkinElmer) was added prior to deployment. The saturating substrate concentrations were determined for each biogeographic province on samples collected at the respective depth. Immediately after filling the polycarbonate bottles, sub-samples (∼100 mL) were transferred from the incubation bottles to the fixation bottles containing 0.2 µm filtered formaldehyde (final concentration 2%) to serve as a killed control (T0), while the live samples were fixed with 2% formaldehyde (final conc.) after 3–12 h of incubation at the respective incubation depth (Extended Data Table 2) at the end of the incubation (Tf) according to the pre-programmed incubation time. All the incubation bottles and tubes in contact with the sample were stored in 0.4–0.5 N HCl overnight, washed three times with MilliQ-water, and rinsed three times with the corresponding 0.2 µm filtered seawater prior to the deployment. The performance of the ISMI has been extensively tested. No significant difference in leucine incorporation was observed between the complete setup of the ISMI and detached ISMI bottles (as a control under atmospheric pressure condition, see below). ^3^H-leucine in the bottles was homogenously distributed as determined in previous tests.

For comparing heterotrophic prokaryotic production under *in situ* pressure with that under atmospheric pressure, water samples were collected at the same depth as the ISMI was deployed using Niskin bottles mounted on a conductivity-temperature-depth (CTD) rosette system (Extended Data Table 1). The hoisting speed of the CTD was 1.0 m s^−1^. Water samples were collected immediately after the CTD arrived on deck of the research vessel and kept in an incubator or water-bath at the respective *in situ* temperature of the sampling depth. The temperature of the water samples collected from the Niskin bottles was typically 2–3°C higher than the *in situ* temperature. Thus, the incubation bottles were incubated for 1–3 h prior to the incubation to attain the *in situ* temperature again (Extended Data Table 2). Sampling of the nepheloid layer was avoided as indicated by the signals of the transmissometer and the optical backscattering sensors mounted on the CTD.

Incubations at atmospheric pressure were performed in identical polycarbonate bottles as used for *in situ* incubations. Three live subsamples and two formaldehyde killed (2% final conc.) controls were used per sample (see Extended Data Table 2) and incubated in temperature-controlled chambers at the same temperature as the *in situ* samples (Extended Data Table 2).

### Bulk heterotrophic prokaryotic biomass production measurements

Leucine incorporation rates were determined according to Kirchman, et al. ^17^. Following formaldehyde fixation of the live samples, samples and controls from *in situ* and atmospheric pressure incubations were filtered onto 0.2 µm polycarbonate filters (25 mm filter diameter, Nuclepore, Whatman). Subsequently, the filters were rinsed twice with 5% ice-cold trichloroacetic acid and twice with Milli-Q water. Filters were air-dried and placed in scintillation vials. Eight mL of scintillation cocktail (either FilterCount or Ultima Gold, PerkinElmer, depending on the research expedition) was added. After about 16 h, the samples were counted in a liquid scintillation counter (Packard, Tri-Carb) on board, and the disintegrations per minute (DPM) obtained were converted into bulk leucine incorporation rates. Additionally, the DPM in 10 µL sample water were determined to check the final concentration of leucine in the incubation vessels of the ISMI.

### MICRO-CARD-FISH

For microautoradiography combined with catalyzed reporter deposition fluorescence *in situ* hybridization (MICRO-CARD-FISH), live samples and formaldehyde-fixed (2% final conc.) controls were incubated at *in situ* and atmospheric pressure conditions as described above. After an incubation time of 3–12 h (Extended Data Table 2), the live samples were fixed with formaldehyde. Upon hoisting the ISMI on board the research vessel, the water contained in the polycarbonate flasks and the samples from the incubations under atmospheric pressure conditions were filtered onto 0.2 µm polycarbonate filters (25 mm filter diameter, GTTP, Millipore) and rinsed twice with Milli-Q water. After drying, the filters were stored at –20°C until further processing. At the home laboratory, the filters were processed as described elsewhere^46^. To permeabilize Archaea, filters were incubated in 0.1 M HCl^47^. Samples were hybridized (at 35°C for 15 h and washing at 37°C for 15 min) with horseradish peroxidase labeled oligonucleotide probes (Extended Data Table 4) and amplified with tyramide-Alexa488 at 46°C for 15 min. After CARD-FISH, the filters were embedded in photographic emulsion (K5, ILFORD) and exposed at 4°C for 14 days in the dark with silica gel as a drying agent. Development and fixing were performed according to the manufacturer’s instructions (developer: Phenisol, ILFORD; fixer: Hypam, ILFORD). Samples were counterstained with 4′,6-diamidino-2-phenylindole (DAPI). Slides were examined on an epifluorescence microscope (Axio Imager M2, Carl Zeiss) equipped with the appropriate filter sets and a camera for photo capturing (≥10 fields). More than 1,000 DAPI-stained cells were enumerated for each CARD-FISH sample. All samples were also hybridized with the antisense probe NON388 (Extended Data Table 4) for unspecific hybridization control. Unspecific binding was always <1% of DAPI-stained cells. Total active cells analysed per sample amounted to: for the mesopelagic cells, *n* ≥ 6,478 at *in situ* and *n* ≥ 6,555 under atmospheric pressure conditions; for bathypelagic cells, *n* ≥ 2,162 at *in situ* and *n* ≥ 1,788 under atmospheric pressure conditions. Cell-specific activity of the different target prokaryotic groups was analyzed by sizing the silver grain halo surrounding probe-positive cells using Axio Vision SE64 Re4.9 (Carl Zeiss). The size of the silver grain area around a cell was converted to single-cell leucine uptake rate (amol leu cell^−1^ d^−1^) based on the regression^19^ obtained using our data set: *R*_halo_ = 9.72 × 10^7^ *R*_leu_ (*r*^2^ = 0.96) where *R*_leu_ is leucine incorporation rate (pmol leu L^−1^ h^−1^) and *R*_halo_ is the total silver grain halo volume (µm^3^ L^−1^ h^−1^) calculated from the area size of the silver grain assuming a spherical distribution. The relatively weak radiation of tritium creates a hemisphere distribution around the cells taking up ^3^H-leucine in the emulsion. Consequently, we calculated the volume of the halo rather than the area (Extended Data Fig. 8). The distribution of cell-specific activities was first expressed as a histogram with a bin interval of 0.17 (amol leu cell^−1^ d^−1^ in log_10_ scale calculated with the smallest number of counts: *n* = 1,788) determined by the kernel estimation based approach^48^. Subsequently, the histogram was used to determine the abundances of piezosensitive, piezotolerant, and piezophilic prokaryotes. Cells with specific activities assigned to the same bin were considered having the same activity. Therefore, when cell-specific uptake rates were classified in the same bin in both *in situ* and atmospheric pressure conditions, these cells were assigned as piezotolerant. Piezosensitive cells were determined as those cells altering their activity from lower to higher activity bins upon depressurization and their minimum and maximum abundances were determined. Accordingly, piezophilic cells were those shifting in the activity bins from higher to lower activity upon depressurization.

### Construction of metagenomic assembled genomes

We used metagenomic assembled genomes (MAGs) to construct a comprehensive gene catalogue for the selected taxa with metagenomic reads using the data set of the Tara Ocean and MALASPINA cruise as well as MAGs from previous publications ^49-51^. The paired-end reads from each metagenome were assembled using MEGAHIT v.1.1.1(k list: 21,29,39,59,79,99,119,141)^52^. The contigs were clustered with two separate automatic binning algorithms: MaxBin^53^ and MetaBAT2^54^ with default settings. The generated genomic bins were de-replicated and refined with Metawrap (bin_refinement). Bins with >70% completeness and <10% contamination (-c 70, -x 10) were kept and pooled with publicly available MAGs^51^ for de-replication using dRep^55^. The phylogenetic affiliation of each MAG was determined using GTDB-Tk^56^. Bacteroidetes-like, Alteromonas-like and SAR202-like MAGs were selected as representatives for downstream analysis. Gene prediction was performed using Prodigal^57^. The predicted genes of each taxa were clustered using 90% similarity applying Cd-hit^58^ to construct a non-redundant protein database, which was used for metaproteomic analysis.

### Metaproteomic analyses of selected bacterial taxa

Metaproteomic data were retrieved from a previous study^28^. Samples for metaproteomic analyses were collected either by Niskin bottles or by *in situ* pumps (WTS-LV, McLane) with 0.2 µm polycarbonate filters mounted^28^. Metaproteomics data were pooled into three groups (epi-, meso-, and bathypelagic) according to depth. The MS/MS spectra from each proteomic sample were searched against the taxa-specific non-redundant protein database using SEQUEST engines^59^ and validated with Percolator in Proteome Discoverer 2.1 (Thermo Fisher Scientific). To reduce the probability of false peptide identification, the target–decoy approach^60^ was used and results <1% false discovery rate (FDR) at the peptide level were kept. Qualified results from peptide-spectrum matches were used for metaproteomic GO enrichment analysis^61^ (metaGOmics, https://www.yeastrc.org/metagomics/home.do) according to the instructions. GO terms with |log_2_ fold change| ≥1 and adjusted *P* value of <0.05 were identified as differentially expressed when comparing samples from different depth layers.

### Calculating the potentially available particulate organic carbon and prokaryotic carbon demand

For estimating the ratio between prokaryotic carbon demand (PCD) and particulate organic carbon (POC) supply, we assembled a large database from published and unpublished prokaryotic ^3^H-leucine incorporation measurements in the Atlantic (*n* = 1,440) and the Pacific (*n* = 783)^13,62-64^. Prokaryotic heterotrophic production (PHP) was calculated using the leucine to carbon conversion factor (CF) of 1.55 kg C mol^−1^ leucine^43^ and 0.44 kg C mol^−1^ leucine^42^. There are higher and lower conversion factors published, however, for our basin-wide production data the applied conversion factors represent the extremes found for specific sites^39^. To calculate prokaryotic heterotrophic production (PHP) rates more typical for *in situ* pressure conditions, we applied the power law fit of Fig. 1 to the measurements performed under atmospheric pressure conditions: PHP_in situ_ = (PHP_atm_ × 494 × *z*^−0.321^)/100; where *z* is depth in meter and PHP_in situ_ and PHP_atm_ are in µmol C m^−3^ d^−1^ under *in situ* and atmospheric pressure conditions, respectively. With these data, the PCD was calculated as PCD = PHP/PGE. From publicly available data a median prokaryotic growth efficiency (PGE) of 8% was applied^42^. A similar value was also reported for the mesopelagic waters in the North Pacific^65^. Consequently, we used a PGE of 8% for mesopelagic depths and a PGE of 3% for bathypelagic waters^13^.

The POC potentially available at a specific depth (POC_a_) was calculated by POC_a_ (mmol m^−3^ d^−1^) = 0.2 × NPP^1.66^ × *z*^−1.68^. The algorithm is based on thorium corrected sediment trap data from the North Atlantic spanning all major biomes^66^, where NPP is the net primary production and *z* is the depth in the water column for which the POC input per day is calculated. The original model calculates fluxes in g C m^−2^ y^−1^, which we converted to mmol C m^−3^ d^−1^ to allow comparing daily rates of PCD with POC input into the specific depth layers. NPP was obtained from the Ocean Productivity website (http://www.science.oregonstate.edu/ocean.productivity) and derived from the Vertically Generalized Production Model (VGPM)^67^ using satellite eight-day averages of chlorophyll. NPP data on the 0.2 × 0.2 degree grid were matched to the nearest degree in longitude and latitude of the stations and the time of sampling for heterotrophic prokaryotic production.

### Analysis and presentation

Statistics and graphics in this study were performed with R version 4.1.1 using RStudio version 1.4.1717 and GMT version 5.4.1. For paired sample tests, normality was checked with Shapiro-Wilk test. If data were normally distributed a *t*-test was performed, otherwise non-parametric tests were applied.

## Acknowledgements

We thank the captain and crew of R/V *Sarmiento de Gamboa*, R/V *Ramon Margalef*, R/V *SONNE*, R/V *Meteor*, R/V *Marion Dufresne* for their support in collecting samples and deployment of the ISMI. We thank M. Varela, C. González-Pola, M. Najdek-Dragić, M. Simon, H. Arndt, I. Obernosterer and J. M. Arrieta for kindly offering opportunities to conduct the study during the cruises and at their laboratories. T. Yokokawa gave advice during the course of the study. We thank M. Álvarez for providing temperature data. B. Mähnert, C. Baranyi, C. Rodríguez, E. Clifford, D. Martinovich helped in instrument preparation, sample collection or processing. Field experiments were conducted during the research cruises: MODUPLAN (CTM-2011-24008), RadProf, SO248 (BacGeoPac), M139 (MerMet 17-97), MOBYDICK, and POSEIDON. This study was supported by JSPS KAKENHI Grant (23651004) to M.U., the Austrian Science Fund (FWF) project I486-B09, Z194 and P28781-B21 to G.J.H. and project P27696-B22 to E.S., the European Research Council under the European Community’s Seventh Framework Program (FP7/2007-2013) / ERC grant agreement No. 268595 (MEDEA project) to G.J.H., and the Austrian Science Fund (FWF) through the PADOM project P23221-B11 to T.R. C.A. was supported by JSPS Postdoctoral Fellowships for Research Abroad (H26–168) and the European Union’s Horizon 2020 research and innovation program under the Marie Sklodowska-Curie No. 701324. The detailed comments of the reviewers helped to improve the overall quality of the manuscript.

## Author contributions

C.A., E.S. and T.R. performed the experiments, analysed leucine incorporation rates, and wrote the manuscript. J.S. helped preparing the ISMI. M.K., J.S. and C.A. analysed the MICRO-CARD-FISH samples. M.U. and C.A. designed the and improved the instrumentation. Z.Z. performed the metagenomics and metaproteomics analyses and wrote the manuscript. G.J.H. designed the study, analysed the data on the carbon budget together with T.R., and contributed to the writing of the manuscript. All authors discussed the results and commented on the manuscript.

## Data sources

Source data are provided in the paper (Extended Data Table 2, Supplementary Table 1 and 2) and are deposited in https://github.com/chie-amano/ismi-data.

## Data availability

Data supporting the findings of this study are available in the paper and its supplementary information files. Station information of the following research cruises is available at the following websites: for the research cruise SO248 (https://doi.org/10.1594/PANGAEA.864673), for M139 (https://doi.org/10.1594/PANGAEA.881298), for MOBYDICK (http://www.obs-vlfr.fr/proof/php/mobydick/mobydick.php).

## Code availability

Computer code is available upon request to the corresponding authors.

## Competing interests

The authors declare no competing interests.

## Extended Data Table

**Extended Data Table 1.** Station details on prokaryotic activity measurements performed in this study. Information is provided for the on-deck atmospheric pressure samples collected by the sampling rosette with the conductivity-temperature-depth (CTD) system.

| Cruise or sampling site | Area | St | Date | Latitude (°) | Longitude (°) | Depth (m) | Temp (°C) | Salinity | DO (μmol kg<sup>-1</sup>) |
| --- | --- | --- | --- | --- | --- | --- | --- | --- | --- |
| MODUPLAN | North Atlantic Ocean | 8 | 5-Aug-2014 | 43.0000 N | 9.7165 W | 445 | 11.39 | 35.67 | 189 |
| MODUPLAN | North Atlantic Ocean | 14 | 8-Aug-2014 | 42.9998 N | 10.6667 W | 1140 | 10.60 | 36.12 | 171 |
| MODUPLAN | North Atlantic Ocean | 108 | 10-Aug-2014 | 43.0000 N | 12.6667 W | 1778 | 4.35 | 35.05 | 233 |
| MODUPLAN | North Atlantic Ocean | 59 | 19-Aug-2014 | 43.9000 N | 3.7833 W | 1974 | 3.91 | 35.03 | 229 |
| MODUPLAN | North Atlantic Ocean | 60 | 20-Aug-2014 | 43.8000 N | 3.7833 W | 990 | 9.72 | 35.78 | 175 |
| MODUPLAN | North Atlantic Ocean | 57 | 21-Aug-2014 | 44.2917 N | 3.7832 W | 1975 | 4.10 | 35.04 | 228 |
| Rovinj* | Adriatic Sea | N/A | 25-Oct-2014 | 45.0856 N | 13.6394 E | 20 | 19.50 | N/D | N/D |
| RadProf15 | North Atlantic Ocean | 11 | 1-Aug-2015 | 43.0253 N | 10.0173 W | 2465 | 3.14 | 34.96 | 220 |
| RadProf15 | North Atlantic Ocean | 111 | 5-Aug-2015 | 43.0010 N | 14.0328 W | 3152 | 2.79 | 34.94 | 213 |
| SO248 | Pacific Ocean | 16† | 26-May-2016 | 49.9999 N | 179.5501 E | 999 | 2.73 | 34.36 | 26 |
| SO248* | Pacific Ocean | 17 | 29-May-2016 | 54.0013 N | 179.5813 E | 501 | 3.63 | 34.08 | 26 |
| M139* | Atlantic Ocean | A1† | 12-Jul-2017 | 15.8860 N | 68.9148 W | 2002 | 4.12 | 34.98 | 197 |
| M139 | Atlantic Ocean | A3† | 22-Jul-2017 | 23.5538 N | 48.0838 W | 3998 | 2.29 | 34.89 | 227 |
| M139 | Atlantic Ocean | A3† | 22-Jul-2017 | 23.5538 N | 48.0838 W | 2000 | 3.68 | 35.00 | 223 |
| M139 | Atlantic Ocean | A5_6† | 31-Jul-2017 | 10.3388 N | 36.9597 W | 3999 | 2.33 | 34.89 | 221 |
| M139 | Atlantic Ocean | A5_6† | 31-Jul-2017 | 10.3388 N | 36.9597 W | 3001 | 2.75 | 34.93 | 225 |
| M139 | Atlantic Ocean | A5_6† | 31-Jul-2017 | 10.3392 N | 36.9617 W | 474 | 8.56 | 34.90 | 69 |
| MOBYDICK | Southern Ocean | M2_1† | 26-Feb-2018 | 50.6160 S | 72.0010 E | 448 | 2.25 | 34.38 | 194 |
| MOBYDICK | Southern Ocean | M3_1† | 3-Mar-2018 | 50.6835 S | 68.0622 E | 1499 | 2.00 | 34.76 | 190 |
| MOBYDICK | Southern Ocean | M2_2† | 6-Mar-2018 | 50.6252 S | 72.0134 E | 400 | 2.16 | 34.30 | 211 |
| MOBYDICK * | Southern Ocean | M1 | 8-Mar-2018 | 49.8502 S | 74.9017 E | 2499 | 1.01 | 34.72 | 197 |
| MOBYDICK | Southern Ocean | M2_3 | 16-Mar-2018 | 50.6159 S | 72.0013 E | 175 | 1.72 | 33.99 | 302 |
| MOBYDICK | Southern Ocean | M3_3 | 18-Mar-2018 | 50.6877 S | 68.0664 E | 1498 | 2.07 | 34.76 | 188 |
| RadProf18 | North Atlantic Ocean | 111 | 24-Aug-2018 | 42.9998 N | 14.0350 W | 3443 | 2.63 | 34.93 | 224 |
| RadProf18 | North Atlantic Ocean | C3 | 27-Aug-2018 | 43.7668 N | 6.1672 W | 743 | 10.23 | 35.67 | 182 |
| POSEIDON | Atlantic Ocean | 5 | 7-Mar-2019 | 36.1575 S | 44.3407 W | 2565 | 2.41 | 34.82 | 204 |
| POSEIDON | Atlantic Ocean | 9 | 12-Mar-2019 | 25.4064 S | 35.0020 W | 3156 | 2.66 | 34.91 | 237 |
| POSEIDON | Atlantic Ocean | 13 | 16-Mar-2019 | 14.4742 S | 30.6714 W | 2962 | 2.69 | 34.90 | 229 |
| POSEIDON | Atlantic Ocean | 19 | 23-Mar-2019 | 5.9091 N | 29.7460 W | 2375 | 3.16 | 34.95 | 229 |
| POSEIDON | Atlantic Ocean | 22 | 26-Mar-2019 | 13.5957 N | 29.7139 W | 3942 | 2.37 | 34.89 | 218 |
| POSEIDON | Atlantic Ocean | 27 | 31-Mar-2019 | 23.7935 N | 23.9188 W | 4002 | 2.40 | 34.90 | 219 |

\*Sample used for instrumental test; †Metaproteomics stations<sup>28</sup>; N/A: not applicable; N/D: not determined; DO- dissolved oxygen measured with an oxygen sensor (Sea-Bird SBE 43 or Aanderaa 4831F).

**Extended Data Table 2.**
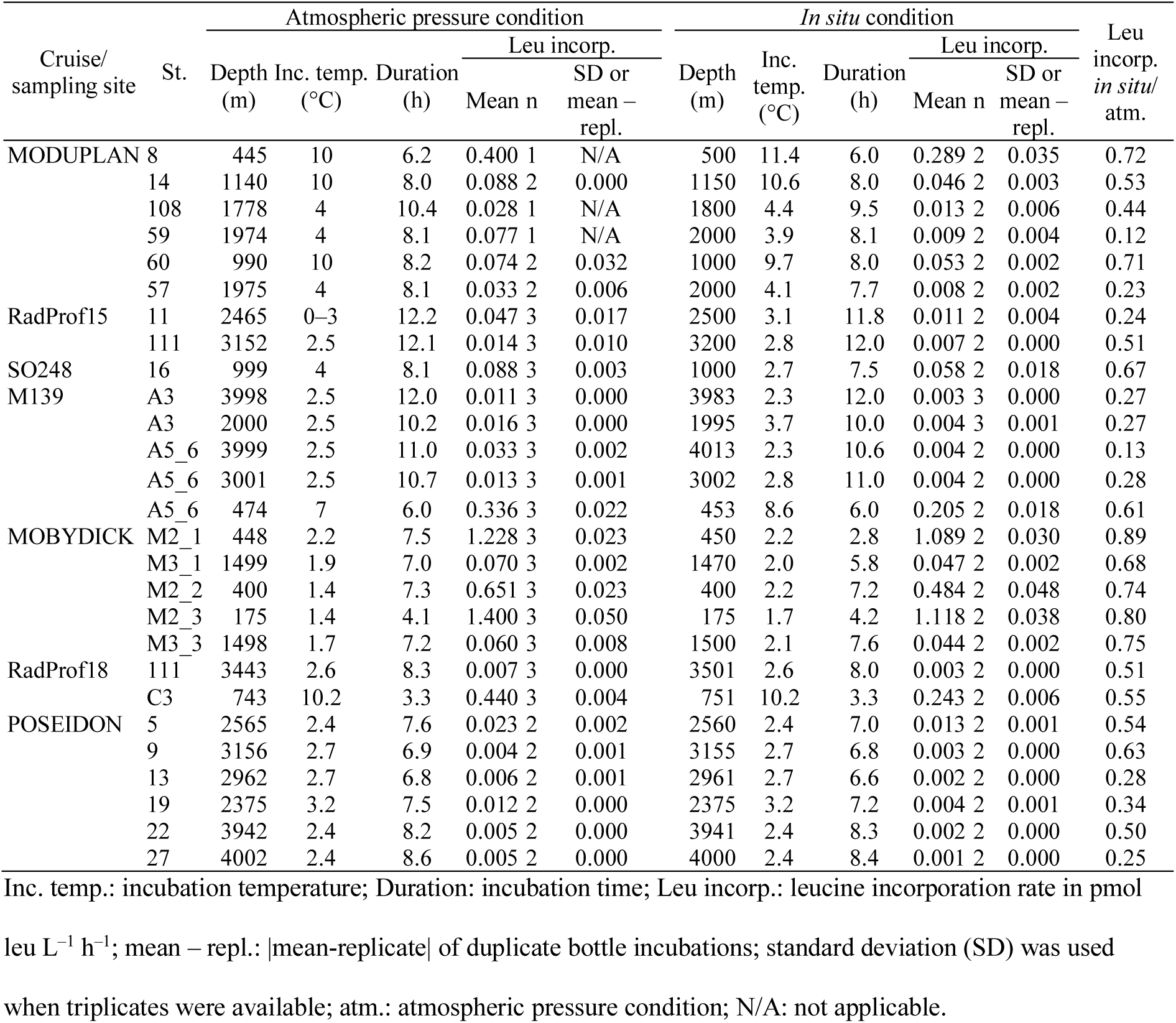
Leucine incorporation rates and incubation conditions under *in situ* and atmospheric pressure conditions.

**Extended Data Table 3.**
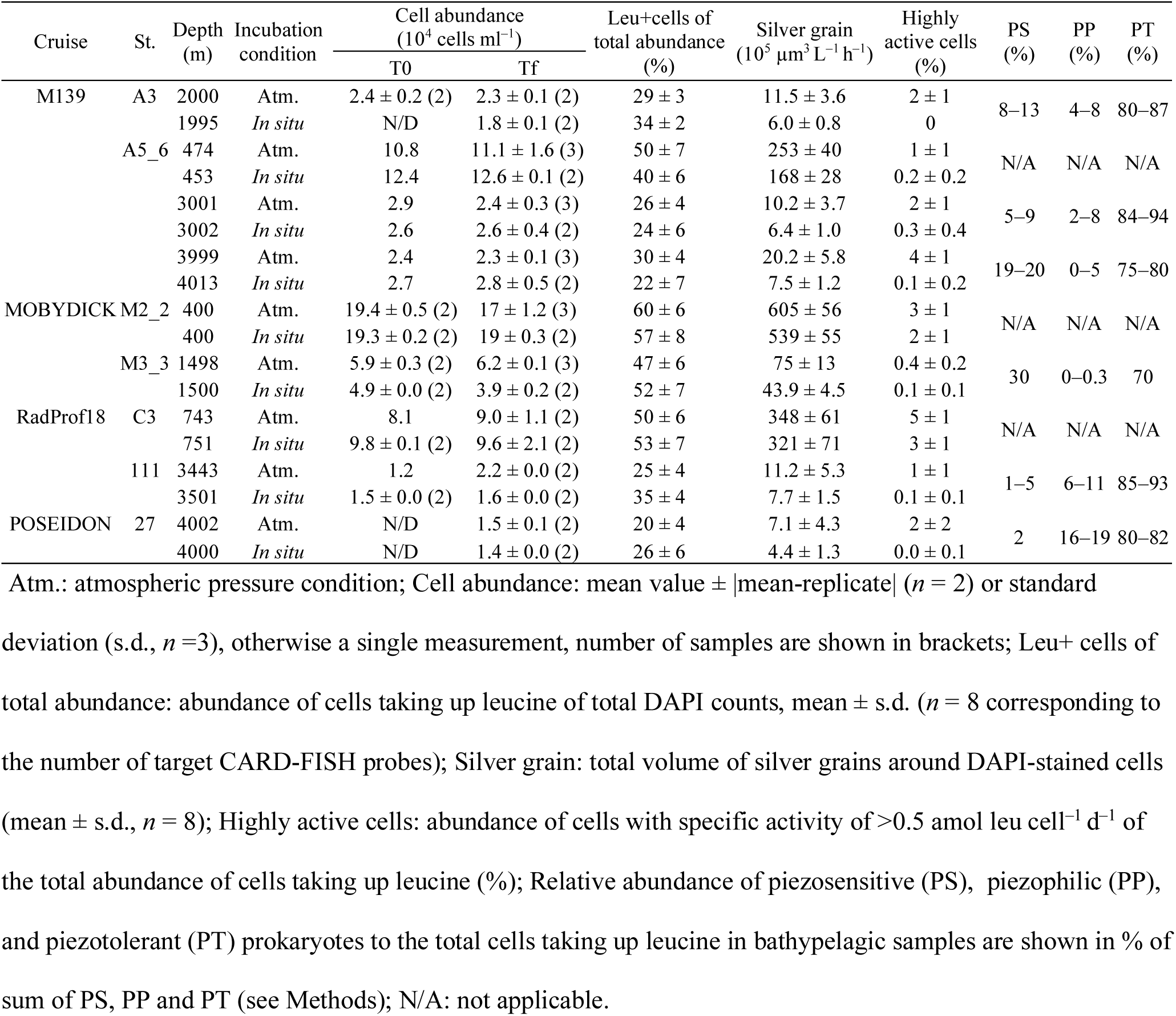
Abundance of cells taking up ^3^H-leucine under *in situ* and atmospheric pressure conditions as determined by MICRO-CARD-FISH.

**Extended Data Table 4.**
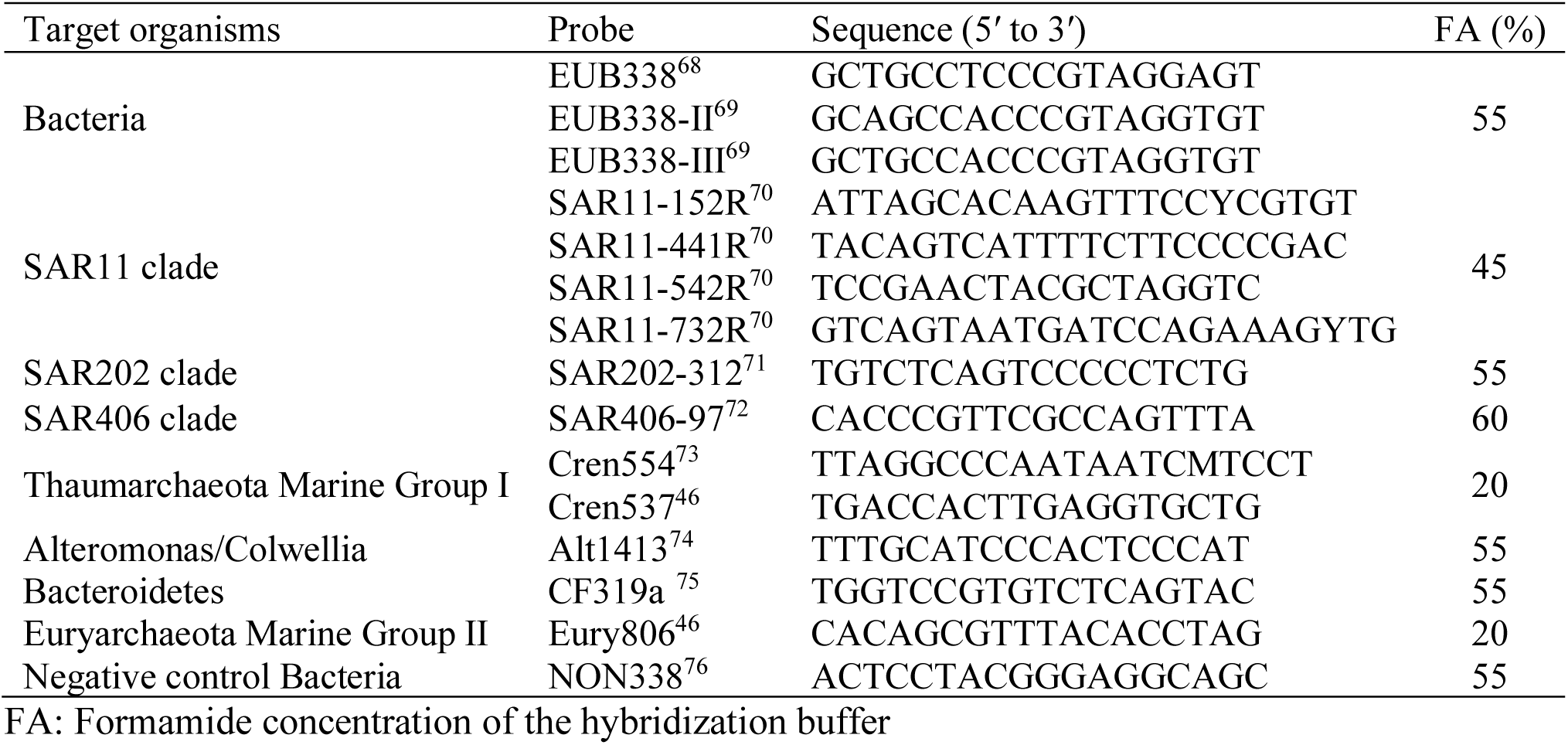
Oligonucelotide probes applied in MICRO-CARD-FISH analyses.

## Extended Data Figure

**Extended Data Fig. 1.**
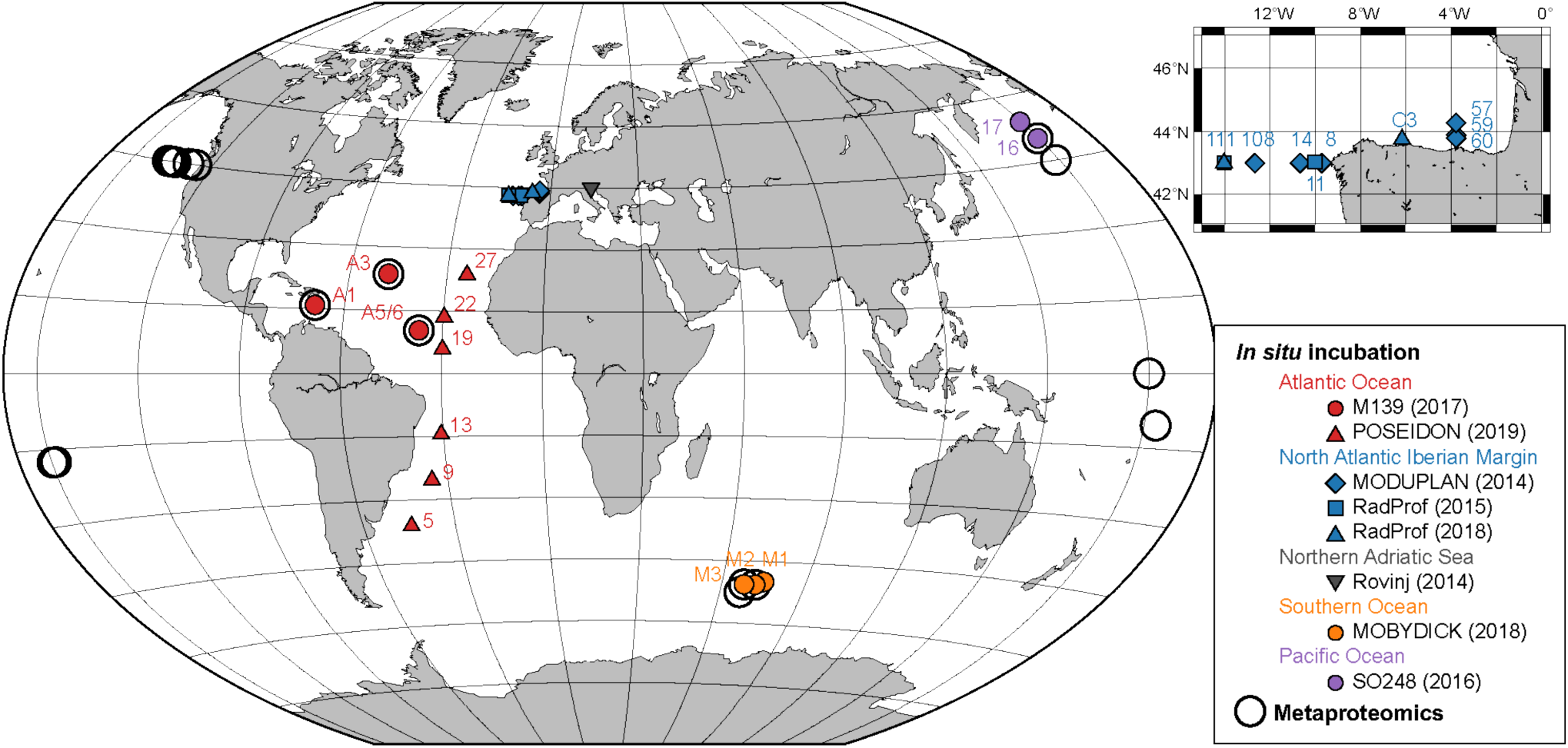
Sampling location of stations where the *in situ* microbial incubator (ISMI) was deployed and metaproteomic analyses were performed. The ISMI was deployed during the M139 and POSEIDON cruise in the Atlantic Ocean, MODUPLAN and RadProf cruises in the North Atlantic off the Iberian Peninsula, MOBYDICK cruise in the Southern Ocean, and SO248 cruise in the Pacific Ocean, and at the Ruđer Bošković Institute, Rovinj, Croatia. Numbers indicate station names. Numbers in brackets indicate the year when sampling was performed. Detailed information of the proteomics stations can be found elsewhere^28^.

**Extended Data Figure 2.**
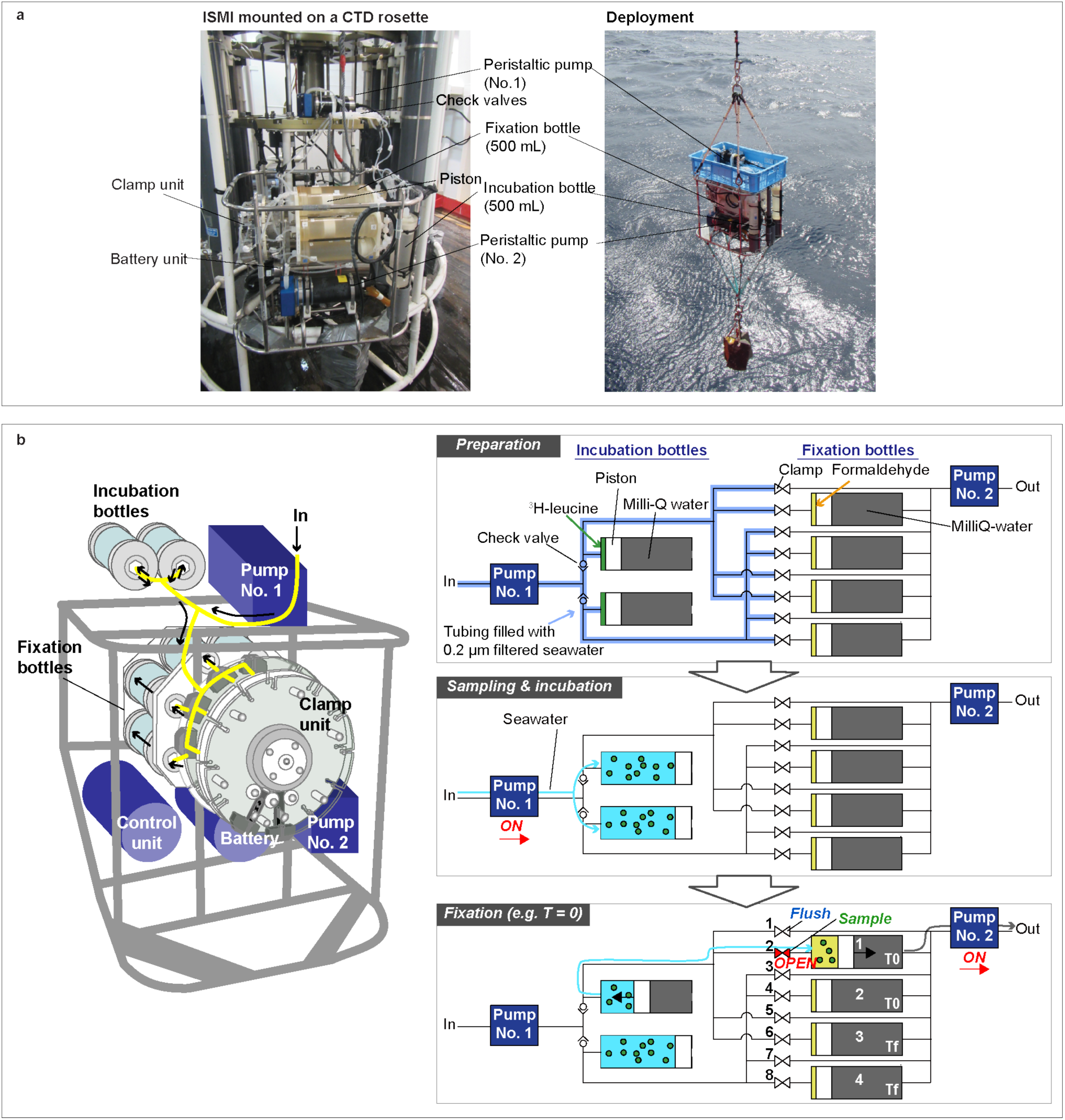
Overview of the ISMI. **a**, The ISMI can be mounted on a rosette sampling system or lowered by the shipboard winch. **b**, Schematic overview of the ISMI. There is only one inlet (left side of the figure) and one outlet (right side) in the system. Prior to deployment, the substrate and the fixative reagent are added into the incubation and fixation cylindrical sampler, respectively. All tubes are pre-filled with either 0.2 µm filtered seawater or MilliQ water. Cylindrical samplers from No. 1 to 4 collect samples in this order by opening the clamps from No. 1 to 8. There is always a flushing step prior to the actual sampling. Incubations are performed either in duplicate or in triplicate.

**Extended Data Fig. 3.**
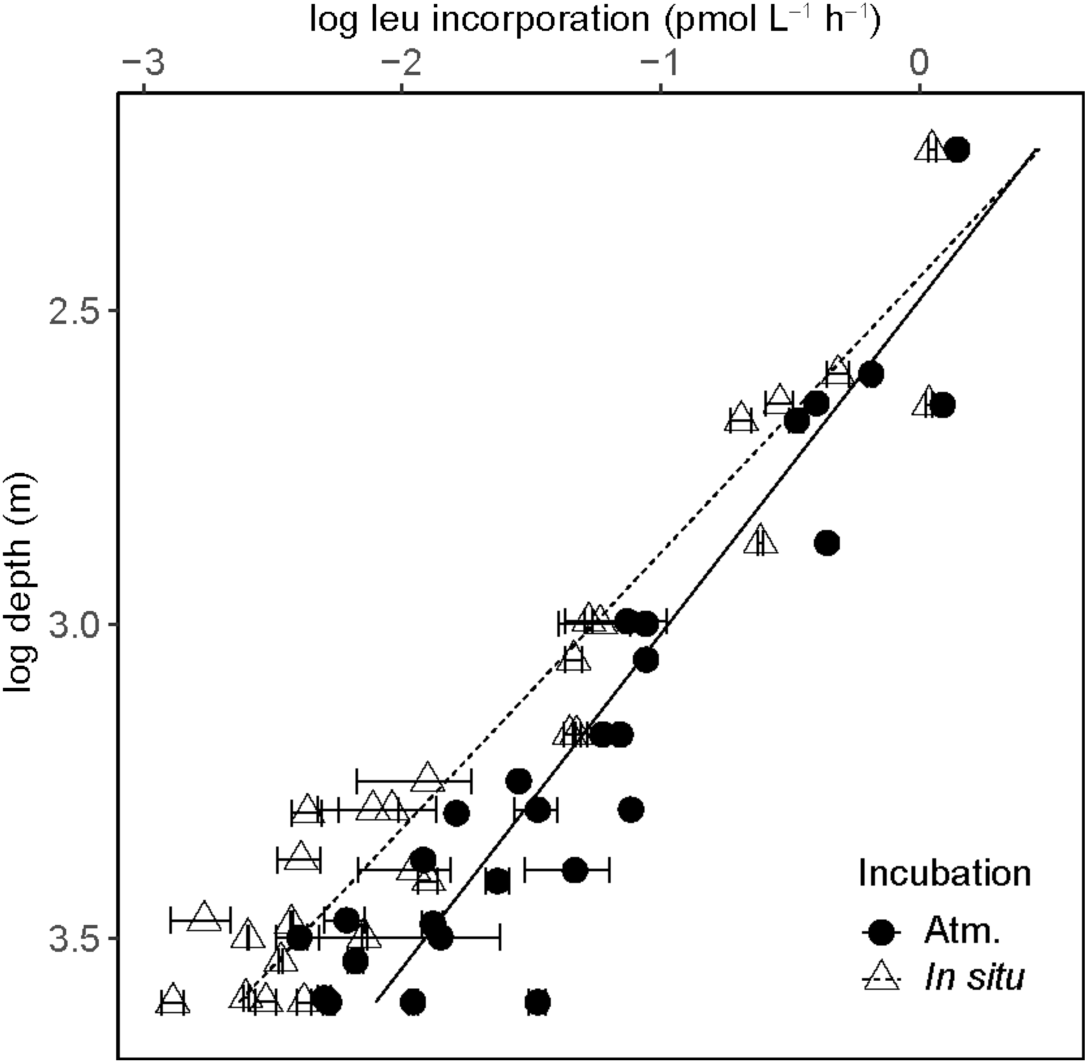
Vertical distribution of leucine incorporation rates incubated under *in situ* and atmospheric pressure conditions. Error bars indicate |mean-replicate| of duplicates (*n* = 2) or s.d. (*n* = 3); Regressions: log (leucine incorporation) (pmol L^−1^ h^−1^) = – 1.9*z* + 4.7 (atm.; *n* = 27, *r*^2^ = 0.87, *P* = 9.9 × 10^−13^); –2.3*z* + 5.6 (*in situ*; *n* = 27, *r*^2^ = 0.92, *P* = 3.9 × 10^−15^) where *z* is log depth in m.

**Extended Data Fig. 4.**
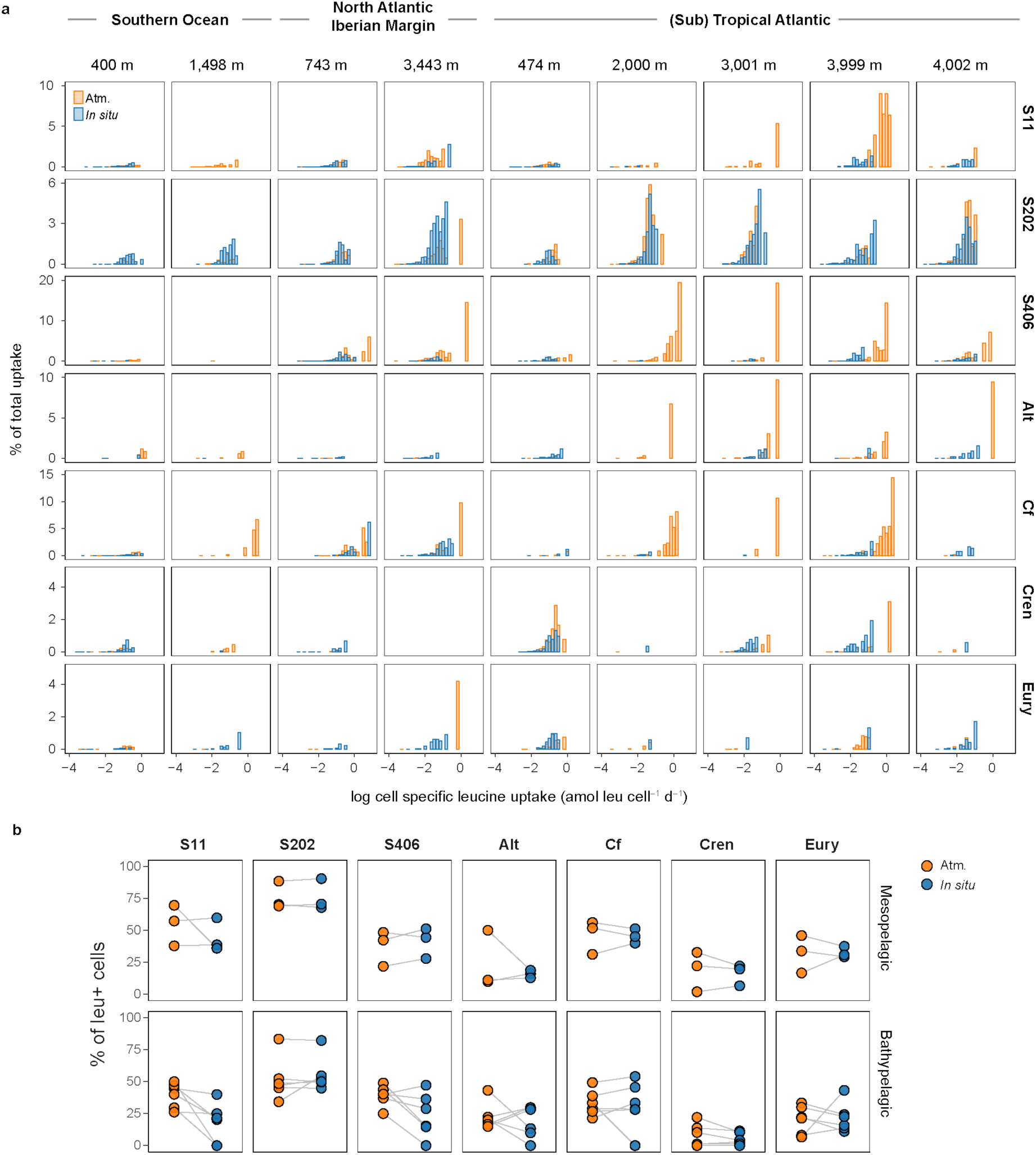
Taxon level response to the hydrostatic pressure. **a**, Cell specific leucine uptake incubated under *in situ* and atmospheric pressure (Atm.) conditions expressed as percentage of total leucine uptake. **b**, Abundance of cells taking up leucine in percent of total abundance of the respective taxon. Target group are indicated as S11: SAR11, S202: SAR202 clade, S406: SAR406 clade, Alt: Alteromonas, Cf: Bacteroidetes, Cren: Thaumarchaeota, Eury: Euryarchaeota.

**Extended Data Fig. 5.**
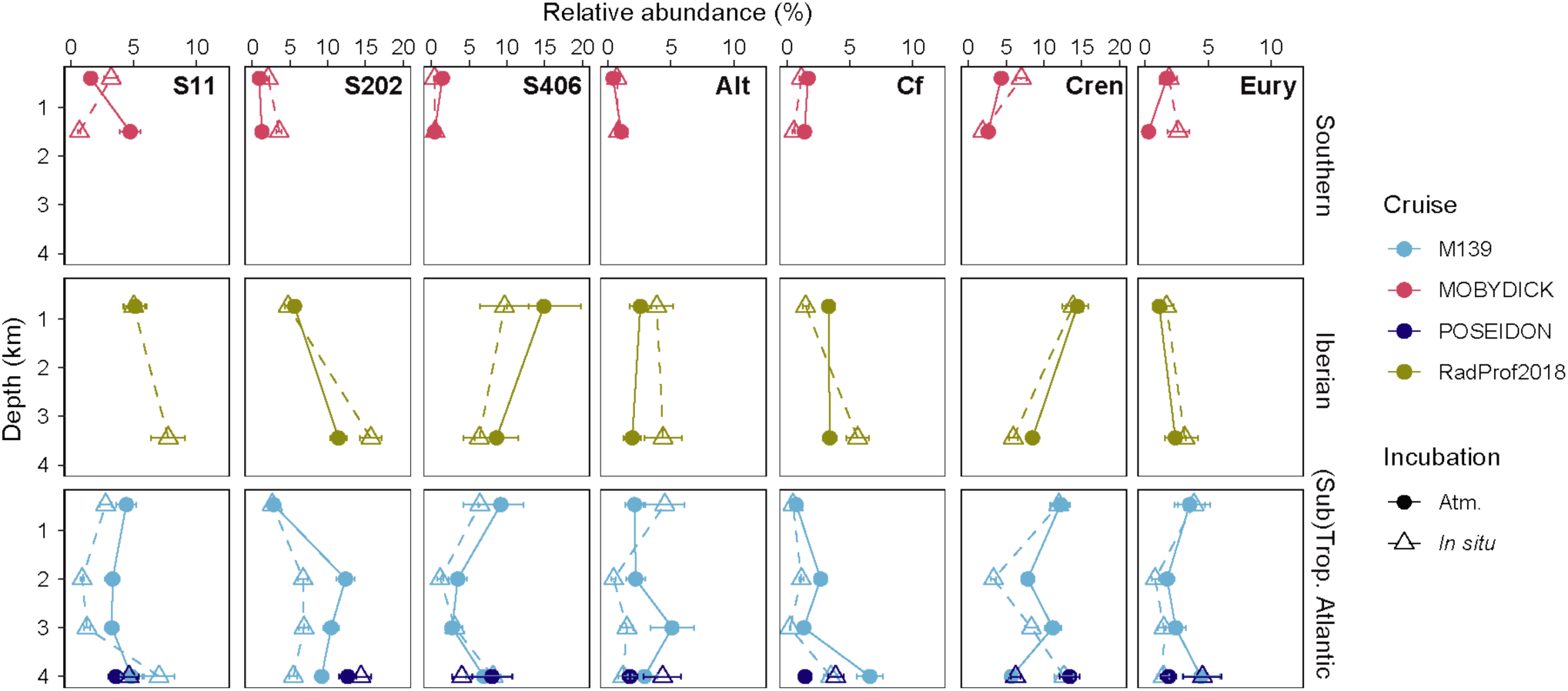
Relative abundance of prokaryotes. Target group are indicated as S11: SAR11 clade, S202: SAR202 clade, S406: SAR406 clade, Alt: Alteromonas, Cf: Bacteroidetes, Cren: Thaumarchaeota, Eury: Euryarchaeota. Error bar shows variations of technical and biological replicates calculated with coefficient of variations (CV). Randomly chosen technical and biological replicates (n ≥ 3) for each taxonomic group were used to calculate the CV of relative abundance. Mean value of the CV was used to estimate the error.

**Extended Data Fig. 6.**
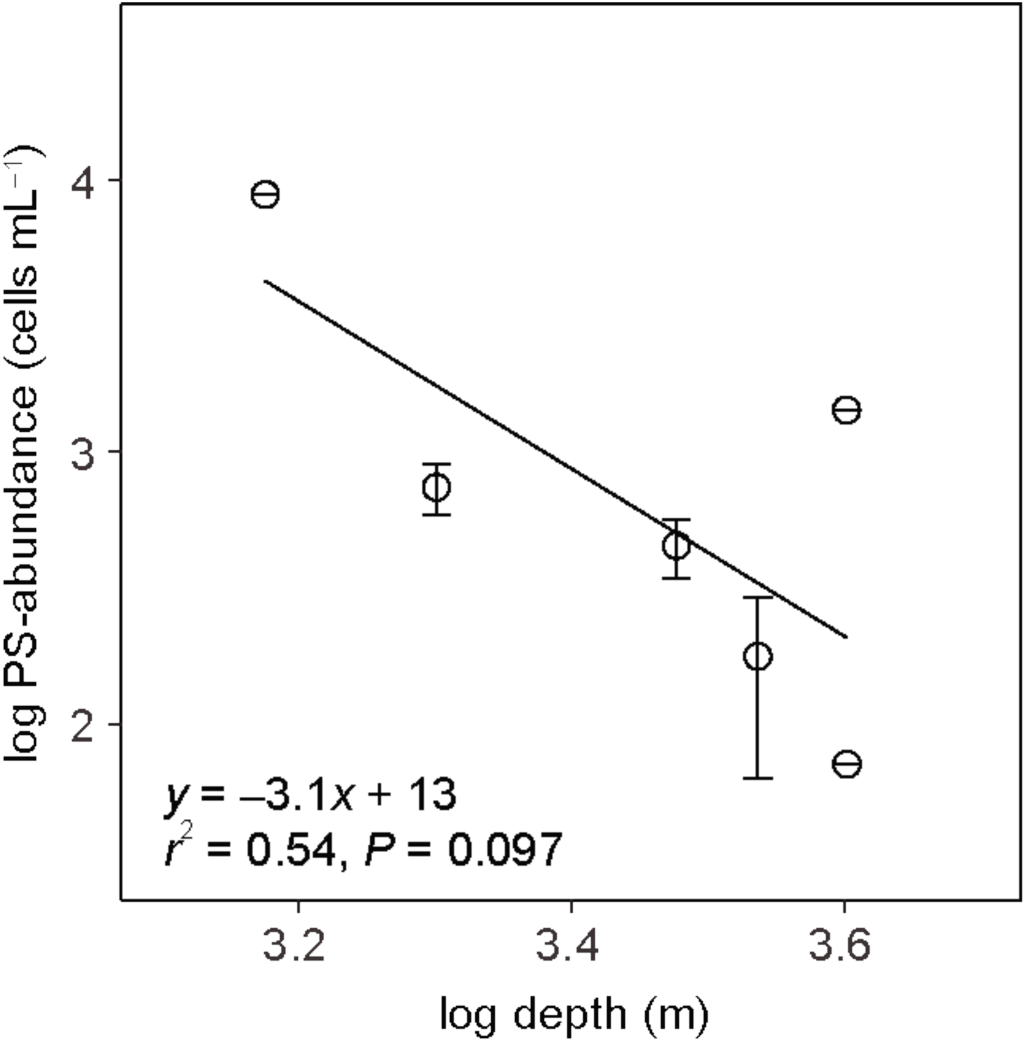
Abundance of piezosensitive microbes (PS) in the bathypelagic waters of the North Atlantic and Southern Ocean. Error bars indicate the minimum and maximum abundance of PS. Mean values indicated by circles were used for calculating the regression.

**Extended Data Fig. 7.**
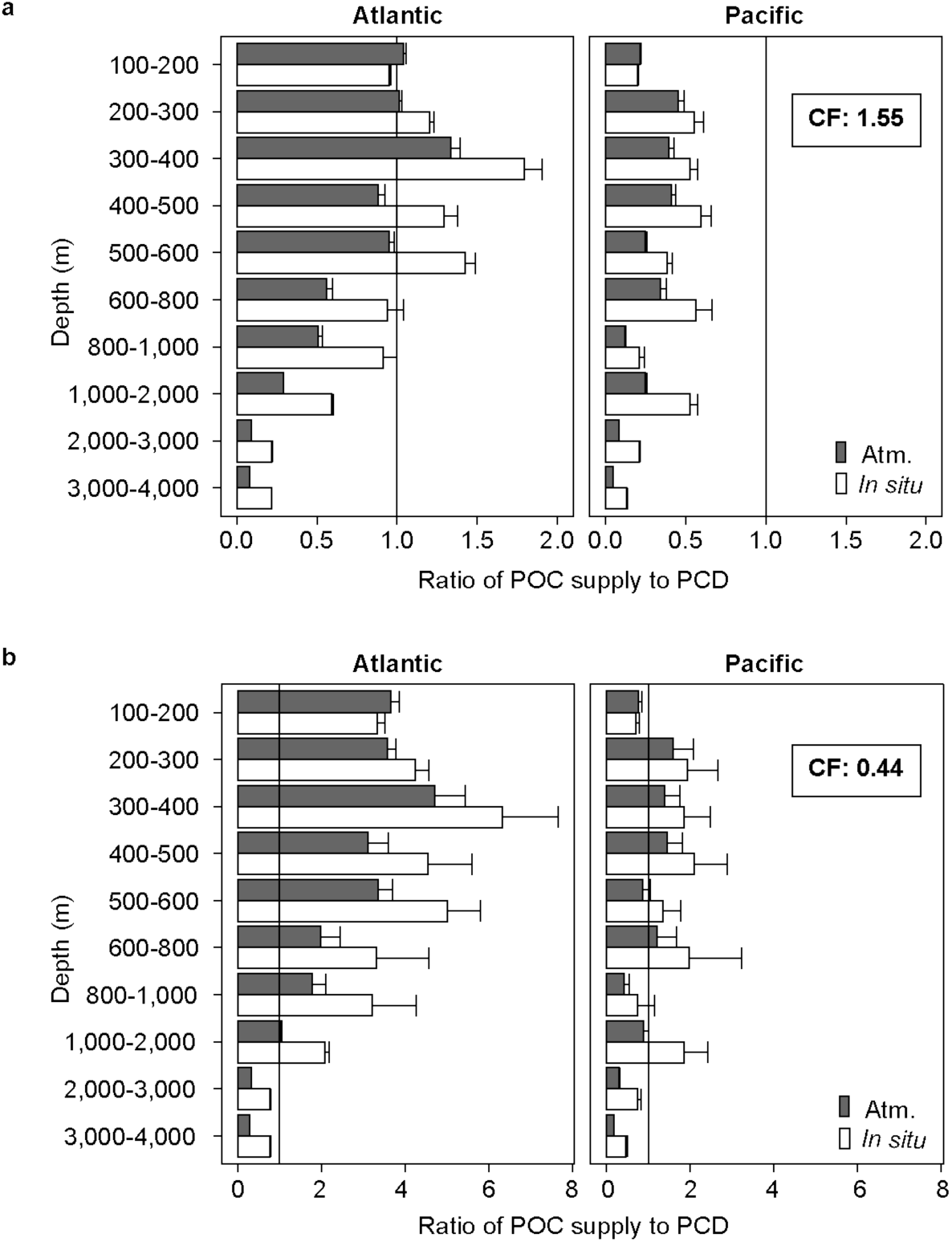
Ratio of modelled particulate organic carbon (POC) supply rate and prokaryotic carbon demand (PCD) calculated from depressurized and *in situ* heterotrophic production rates in the Atlantic and the Pacific Ocean. The particulate organic carbon (POC) potentially available at a specific depth is calculated using depth dependent sediment trap data^66^ and satellite derived net primary production estimates. The prokaryotic carbon demand assumes a grand average of 8% growth efficiency for the meso- and 3% for the bathypelagic waters. PCD was calculated using leucine to carbon conversion factors of **a**, 1.55 kg C mol^−1^ leu and **b**, 0.44 kg C mol^−1^ leu (see Methods). A ratio of 1 indicates that the POC supply rate matches PCD. Values <1 suggest inadequate supply of POC to support the PCD. Error bars indicate standards errors taking error propagation into account.

**Extended Data Fig. 8.**
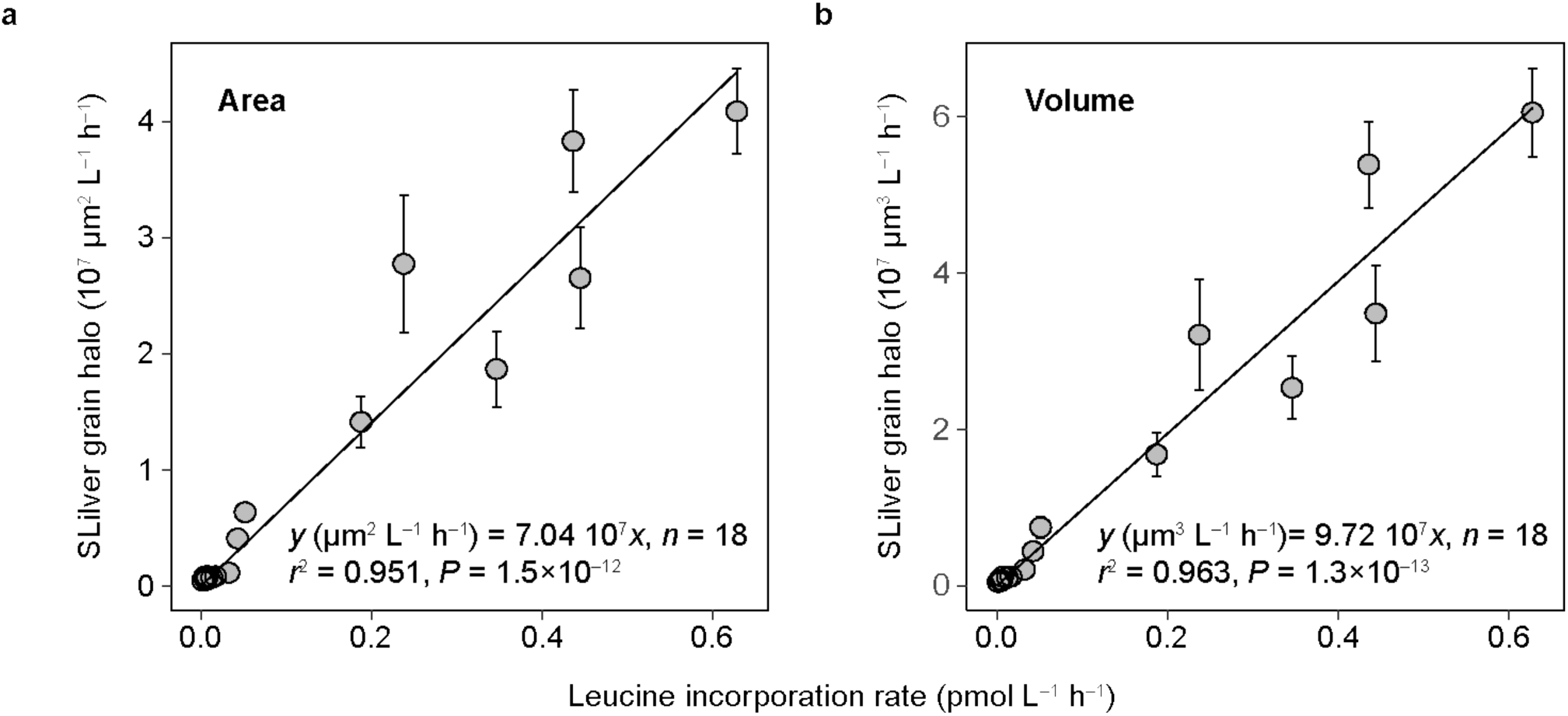
Single-cell activity measurements. Relationship of bulk leucine incorporation rates and silver grain halo around cells expressed as **a**, area and **b**, volume. All the MICRO-CARD-FISH samples under *in situ* and atmospheric pressure conditions at 9 stations (see Methods) were used to obtain the regressions. Error bars are standard deviations of 8 technical replicates which correspond to the number of probes used. A slightly better correlation between bulk leucine incorporation and silver grain halo volume than with halo area was found. Consequently, single-cell leucine uptake was calculated using the regression with halo volume in this study.

**Supplementary Table 1**. List of up-regulated functions of Alteromonas, Bacteroidetes and SAR202 using gene ontology (up-regulation.csv).

**Supplementary Table 2**. List of down-regulated functions of Alteromonas, Bacteroidetes and SAR202 using gene ontology (down-regulation.csv).

## Notes

### Competing Interest Statement

The authors have declared no competing interest.

